# The UCSC Xena platform for public and private cancer genomics data visualization and interpretation

**DOI:** 10.1101/326470

**Authors:** Mary Goldman, Brian Craft, Mim Hastie, Kristupas Repečka, Fran McDade, Akhil Kamath, Ayan Banerjee, Yunhai Luo, Dave Rogers, Angela N. Brooks, Jingchun Zhu, David Haussler

**Affiliations:** UC Santa Cruz Genomics Institute, University of California, Santa Cruz, CA, USA; Clever Canary, New York, NY, USA; Vilnius University, Vilnius, Lithuania; Birla Institute of Technology and Science, Goa, India; National Institute of Technology, Durgapur, India; Department of Genetics, Stanford University, Stanford, CA USA; Department of Biomolecular Engineering, University of California, Santa Cruz, CA, USA

## Abstract

UCSC Xena is a visual exploration resource for both public and private omics data, supported through the web-based Xena Browser and multiple turn-key Xena Hubs. This unique archecture allows researchers to view their own data securely, using private Xena Hubs, simultaneously visualizing large public cancer genomics datasets, including TCGA and the GDC. Data integration occurs only within the Xena Browser, keeping private data private. Xena supports virtually any functional genomics data, including SNVs, INDELs, large structural variants, CNV, expression, DNA methylation, ATAC-seq signals, and phenotypic annotations. Browser features include the Visual Spreadsheet, survival analyses, powerful filtering and subgrouping, statistical analyses, genomic signatures, and bookmarks. Xena differentiates itself from other genomics tools, including its predecessor, the UCSC Cancer Genomics Browser, by its ability to easily and securely view public and private data, its high performance, its broad data type support, and many unique features.

## Introduction

There is a great need for easy-to-use cancer genomics visualization tools for both large public data resources such as TCGA (The Cancer Genome Atlas) (Chin 2011) and the GDC (Genomic Data Commons) (Grossman 2016), as well as smaller-scale datasets generated by individual labs. Commonly used interactive visualization tools are either web-based portals or desktop applications. Data portals have a dedicated backend and are a powerful means of viewing centrally hosted resource datasets (e.g. Xena’s predecessor, the UCSC Cancer Browser (currently retired, Zhu 2009), cBioPortal (Cerami 2012), ICGC (International Cancer Genomics Consortium) Data Portal (Zhang 2019), GDC Data Portal (Grossman 2016)). However, researchers wishing to use a data portal to explore their own data have to either a redeploy the entire platform, a difficult task even for bioinformaticians, or upload private data to a server outside the user’s control, a non-starter for protected patient data such as germline variants (e.g. MAGI (Mutation Annotation and Genome Interpretation, Leiserson 2015), WebMeV (Wang 2017), Ordino (Streit 2018)). Desktop tools can view a user’s own data securely (e.g. IGV (Integrated Genomics Viewer, Thorvaldsdóttir 2013), Gitools (Perez-Llamas 2011)), but lack well-maintained, prebuilt files for the ever-evolving and expanding public data resources. This dichotomy between data portals and desktop tools highlights the challenge of using a single platform for both large public data and smaller-scale datasets generated by individual labs.

Complicating this dichotomy is the expanding amount, and complexity of, cancer genomics data resulting from numerous technological advances, including lower-cost high-throughput sequencing and single-cell based technologies. Cancer genomics datasets are now being generated using new assays such as whole-genome sequencing (Campbell 2017), DNA methylation whole-genome bisulfite sequencing (Zhou 2018), and ATAC-seq (Assay for Transposase-Accessible Chromatin using sequencing, Corces 2018). Visualizing and exploring these diverse data modalities is important but challenging, especially as many tools have traditionally specialized in only one or perhaps a few data types. And while these complex datasets generate insights individually, integration with other –omics datasets is crucial to help researchers discover and validate findings.

UCSC Xena was developed as a high-performance visualization and analysis tool for both large public repositories and private datasets. It was built to scale with the current, and future, data growth and complexity. Xena’s unique privacy-aware architecture enables cancer researchers of all computational backgrounds to explore large diverse datasets. Researchers use the same system to securely explore their own data, together or separately from the public data, all while keeping private data secure. The system easily supports many tens of thousands of samples and has been tested up to as many as a million cells. The simple and flexible architecture supports a variety of common and uncommon data types. Xena’s unique Visual Spreadsheet visualization integrates gene-centric and genomic-coordinate-centric views across multiple data modalities, providing a deep, comprehensive view of genomic events within a cohort of tumors.

### Xena’s privacy-aware architecture

UCSC Xena (http://xena.ucsc.edu) has two components: the frontend Xena Browser and the backend Xena Hubs (Figure 1). The web-based Xena Browser empowers biologists to explore data across multiple Xena Hubs with a variety of visualizations and analyses. The backend Xena Hubs host genomics data from laptops, public servers, behind a firewall, or in the cloud, and are configured to be public or private (Supplemental Figure 1). The Xena Browser receives data simultaneously from multiple Xena Hubs and integrates them into a single coherent visualization within the browser.

**Figure 1.**
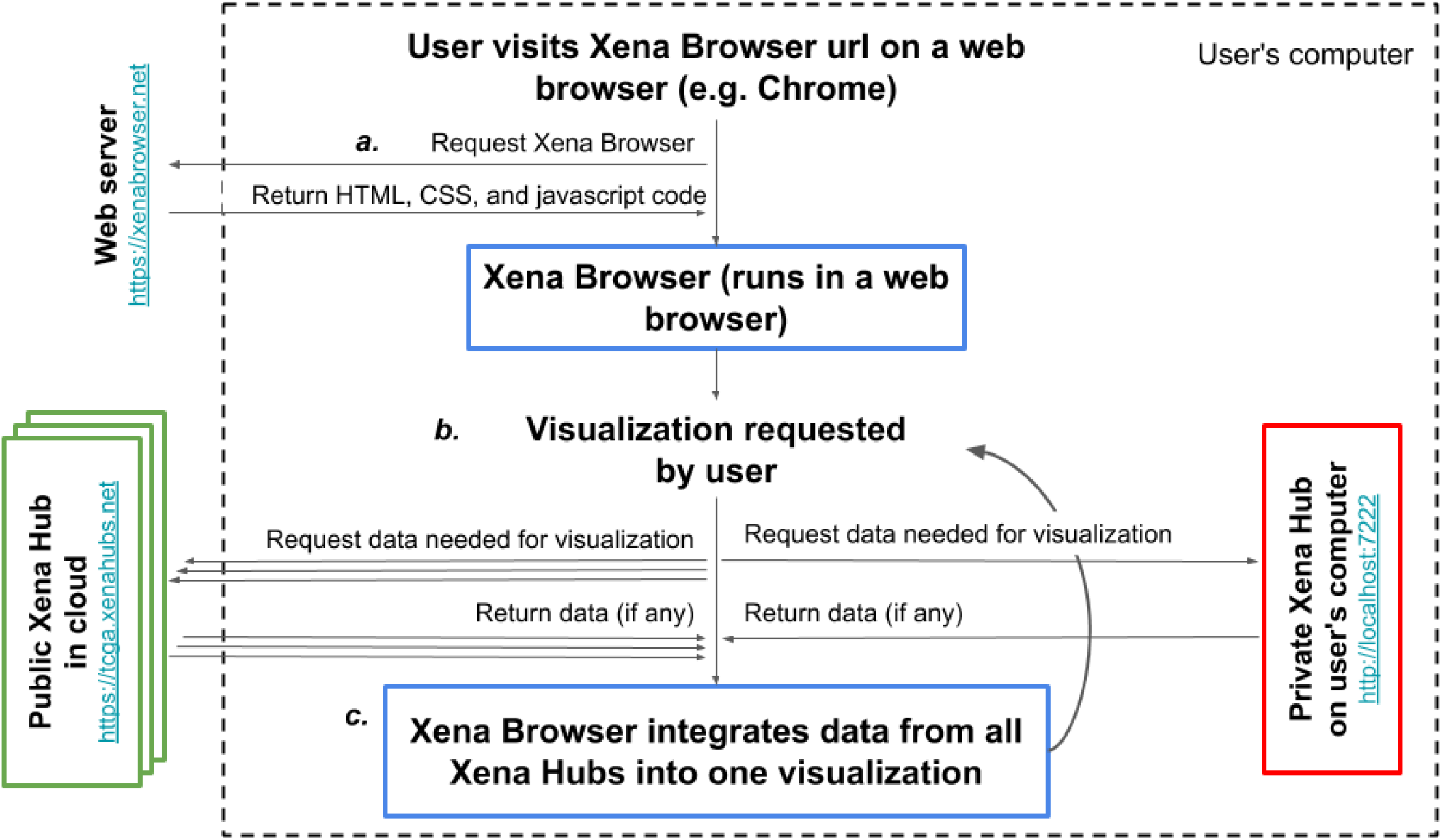
Xena’s architecture to securely join public and private data. Data always flows from the Xena Hubs to the Xena Browser for visualization and integration. **a**) User’s web browser (e.g. Chrome) requests the Xena Browser code and runs it. **b**) Using the Xena Browser, the user requests a visualization, initiating a request for data from the Xena Browser’s list of public hubs. Simultaneous with this request, the Xena Browser requests data from the private local hub on the user’s computer. **c**) The Xena Browser code combines data from all Xena Hubs together into one coherent visualization. The user can then interact with the visualization to trigger a new data request.

There are two types of private Xena hubs (Supplemental Figure 2). The first type is a hub installed on a user’s own computer. It is configured to only respond to requests from the computer’s localhost network interface (i.e. http://127.0.0.1). This ensures that the hub only communicates with the computer on which the hub is installed. The second type of private hub is one configured to respond to requests from external computers, however access to the computer is controlled via a firewall or similar technology. The hub uses the security provided by the computer to secure the data. This model takes advantage of security that is typically already in place to protect such data, thereby reducing user workload by not requiring re-authorization. Users who host this type of private data will share the URL with authorized individuals. Other users who may inadvertently acquire knowledge of the protected hub will not be able to connect due to the firewall. This second type of Xena Hub is useful to share private data within a lab or institution.

In addition to the private hubs, there are public Xena hubs, that is hubs that are configured to respond to requests from external computers and are not blocked by a firewall (Supplemental Figure 2). Public hubs enable data sharing by hosting large public resources. While we host a number of public hubs (Supplemental Table 1), users can also set up their own. An example of one is the Treehouse Hub set up by the Childhood Cancer Initiative to share pediatric cancer RNA-seq gene expression data (Supplemental Note).

Public and private Xena Hubs use the same software; the only difference is in their configuration. Hubs default to only respond to requests from the computer’s localhost network, locking down data accessibility to the host computer. Hubs only respond to external network requests if a user configures the hub to do so. Xena Hubs are designed to be turn-key, allowing users who may not be computationally savvy to easily install and use a Xena Hub on their personal computer (https://xena.ucsc.edu/private-hubs/). An interactive setup wizard guides users through the process of installing and running a Xena Hub on their Windows or Mac computers, while a web wizard guides them through the data loading process (https://tinyurl.com/localXenaHub). In addition to these user-friendly wizards, Xena Hubs can be installed and used via the command line on Windows, Mac and Linux machines.

The Xena Browser automatically connects to a default list of the public hubs we host (Supplemental Table 1), and, if it exists, the private local hub on users’ computer. Users can add a new public or private hub by entering the hub URL address on the Xena Browser on the Data Hubs page. Data integration occurs only within the Xena Browser, keeping private data secure. Genomic data flows from a Xena Hub to the Xena Browser (Figure 1), and never communicates the data it displays back to any server. The only exception to this model occurs when saving bookmarks as URLs, a feature that allows users to save live views of their current visualization. If a visualization contains only data from public Xena Hubs, users can generate a URL for their current view, which will take researchers back to the live browser session. Since the data is already public, we store the data in view for each URL on our web server, allowing it to be shared with colleagues or included in presentations. If a view contains any data from a non-public Xena Hub, users are instead required to download the current visualization as a file. By giving users a file instead of a URL, we ensure that we never keep user’s private data on our public servers. This file can then be shared via email, etc and then imported back into the Xena Browser to recreate the live browser session. Thus, even when bookmarking, protected data is kept secure through Xena’s architecture and the use of private hubs.

In addition to its security advantages, Xena’s unique architecture of a decoupled Xena Browser and multiple Xena Hubs enables several other features. First, researchers can easily view their own private data by installing their own Xena Hub. Xena Hubs are lightweight compared to a full-fledged application and install easily on most computers. Second, users can use the same platform to view both public and private data together. Xena integrates data across multiple hubs, allowing users to view data from separate hubs as a coherent data resource (Figure 2). Xena does this while keeping private data secure and avoiding the need to download large public resources. This is especially useful for researchers who wish to view their own analysis results on public data, such as their own clustering calls, but don’t want to host a separate version of these resources. Third, the Xena platform scales easily. As more datasets are generated, more Xena Hubs are added to the network, effectively growing with expanding genomics resources. These advantages, in addition to the security advantages, are a major departure from and innovation over the UCSC Cancer Browser.

**Figure 2.**
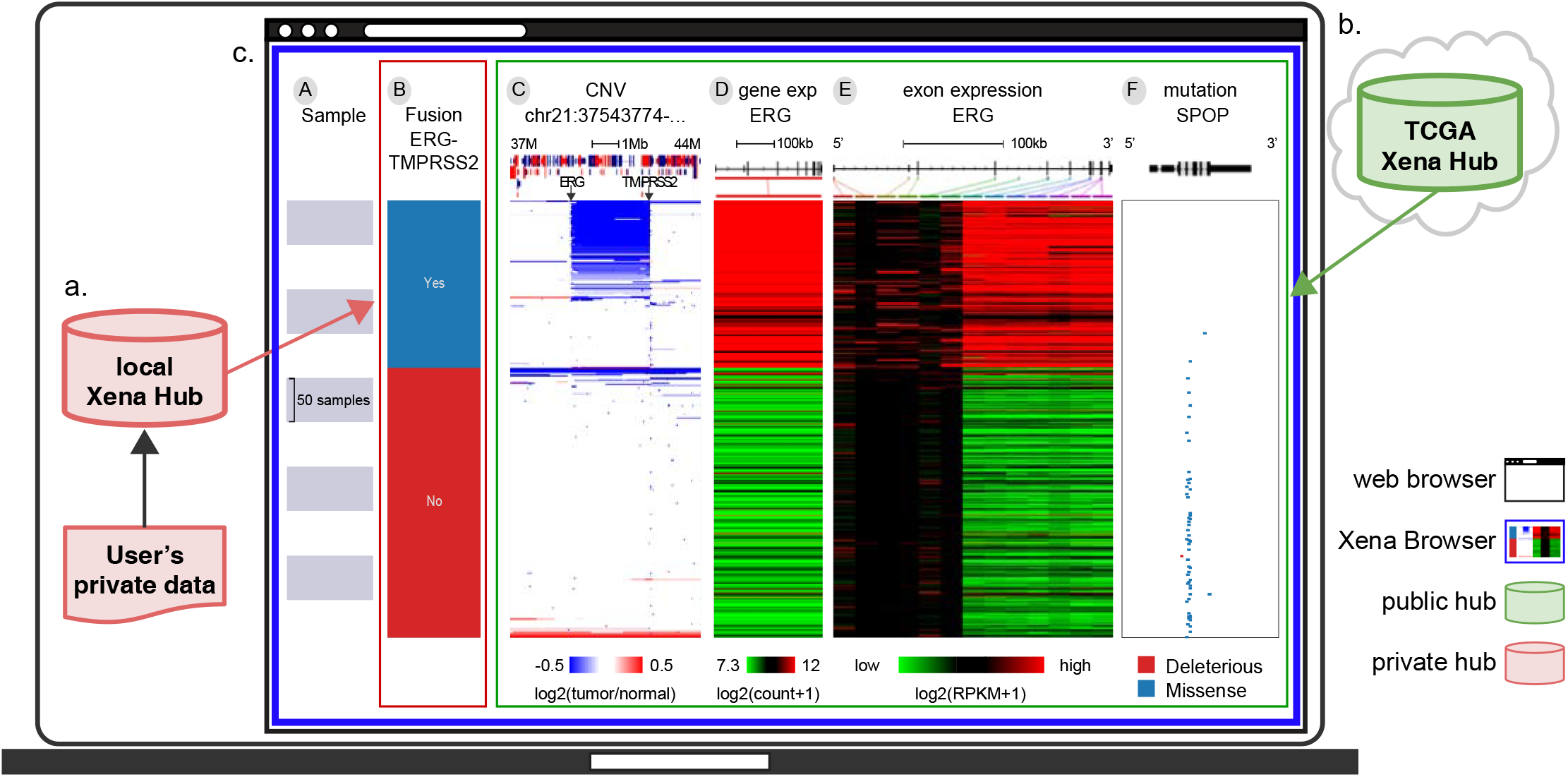
An example Xena Browser Visual Spreadsheet examining published ERG (ETS Transcription Factor) - TMPRSS2 (Transmembrane Serine Protease 2) fusion calls in TCGA PRAD (prostate cancer) by combining data from local and public Xena Hubs together. **a)** A user downloaded ERG-TMPRSS2 fusion calls on TCGA PRAD samples from Gao et al. 2018 (n=492) and loaded the data into their own local Xena Hub. **b)** TCGA copy number, gene expression and mutation data from the same samples are available via the public TCGA hub. **c)** The user then compared the fusion calls to the public data using Xena Browser Visual Spreadsheet. Column B is the fusion call from Gao et al. Column C is copy number variation data, zoomed in to a region of chromosome 21 (37-44Mb). Amplifications are in red and deletions are in blue. The diagram at the top shows genes along the chromosome, where red genes are on the positive strand and blue are on the negative strand. Columns D is ERG gene expression and Column E is ERG exon expression. Expression is colored red to green for high to low expression. The gene diagram at the top shows exons as boxes, with tall coding regions and shorter untranslated regions. Column F is SPOP (Speckle Type BTB/POZ Protein) mutation status and also has a gene diagram at the top. The position of each mutation is marked in relation to the gene diagram and colored by its functional impact: deleterious mutations are red and missenses are blue. We can see that the fusion calls are highly consistent with the characteristic overexpression of ERG (columns D, E). However, only a subset of those samples in which a fusion was called can be seen to also have the fusion event observed in the copy number data via an intra-chromosomal deletion of chromosome 21 that fuses TMPRSS2 to ERG as shown in column C. This observation is consistent with the 63.3% validation rate described in Gao et. al. 2018. SPOP mutations (blue tick marks in column F) are mutually exclusive with the fusion event. Rows are sorted by the left-most data column (column B) and subsorted on columns thereafter.

#### Xena Browser visualizations and functionalities

The Xena Browser (https://xenabrowser.net) has a wide variety of visualizations and analyses including survival analyses, scatter plots, bar graphs, statistical tests, genomic signatures, as well as our unique Visual Spreadsheet view. The Xena Visual Spreadsheet was designed to enable and enhance integration across diverse data modalities, providing researchers a more biologically complete understanding of genomic events and tumor biology. Analogous to an office spreadsheet application, it is a visual representation of a data grid where each column is a slice of genomic or phenotypic data (e.g. gene expression, mutation calls, methylation probes, subtype classifications, or age), and each row is a single entity (e.g. a bulk tumor sample, cell line, or single cell) (Figure 2). Xena’s Visual Spreadsheet displays genomic data in wide variety of gene-centric, coordinate-centric, and feature-centric views (Supplemental Figure 3) for both coding and non-coding regions (Supplemental Figure 4). Dynamic web links to the UCSC Genome Browser give genomic context to any gene, chromosome region, or feature. Researchers can easily re-order the Visual Spreadsheet, hierarchically cluster genes, zoom in to just a few samples or out to the whole cohort, all leading to an infinite variety of views in real time. These dynamic views enable the discovery of patterns among genomic and phenotype parameters, even when the data are hosted across multiple data hubs (Figure 2).

The power of the Visual Spreadsheet is its deep data integration. Integration across different data modalities, such as gene expression, copy number variation, and DNA methylation, gives users a more comprehensive view of a genomic event in a tumor sample. For example, Xena’s Visual Spreadsheet can help elucidate if higher expression for a gene is driven by copy number amplification, or by a missense mutation (Supplemental Figure 5), or by demethylation and opening of the promoter region as reflected in the DNA methylation and ATAC-seq data (Supplemental Figure 3). Integration across gene- and coordinate-centric views helps users examine genomic events in different chromosome contexts. For example, Xena’s Visual Spreadsheet can help elucidate if a gene amplification is part of a chromosomal arm duplication or a focal amplification (Supplemental Figure 6). Integration across genomic and clinical data gives users the ability to make connections between genomic patterns and clinically relevant phenotypes such as subtypes. For example, Xena’s Visual Spreadsheet can help elucidate if increased HRD (Homologous Recombination Deficiency) signature scores are enriched in a specific cancer type or subtypes (Supplemental Figure 7). Finally, integration across user’s own data and public resources on the same samples helps users to gain insights into their own data. For example, Xena’s Visual Spreadsheet can help a researcher see how a fusion call from the literature relates to the expression of other downstream genes (Figure 2). By not differentiating between public data and private data from rendering perspective, it appears to the user that all data come from a coherent source. These diverse integrations help researchers harness the power of comprehensive genomics studies, either their own or of public resources, driving discovery and a deeper understanding of cancer biology.

In addition to the Visual Spreadsheet, Xena has many additional powerful views, analyses, and functionalities. Our powerful text-based search allows users to dynamically highlight, filter, and group samples (Supplemental Figure 8). Researchers use this to search the data on the screen similar to the ‘find’ functionality in Microsoft Word. Samples are matched and highlighted in real-time as the user types. Researchers can then filter to their samples of interest, or dynamically build subgroups.

This is a powerful way to dynamically construct sub-populations based on any genomic data for comparison and analysis. Xena also has highly configurable Kaplan-Meier analyses, bar charts, box plots, and scatter plots, all with statistical tests automatically computed (Supplemental Figure 7, Supplemental Figure 9). We support data sharing through bookmarked views and high resolution PDFs. Genomic signatures are easily built over gene expression data or any other genomic data type.

Performance is critical for interactive visualization tools, especially on the web. Growing sample sizes for genomic experiments has become a challenge for many tools, including for the UCSC Cancer Browser. Knowing this, we optimized Xena to support visualizations on many tens of thousands of samples, delivering slices of data in milliseconds to a few seconds. During the first 8 months in 2019, we averaged 1,400 users/week with an average concurrent current usage of 3.34 users. To ensure we will continue to be performant as we scale, we tested our public hubs deployed in the cloud with 50 concurrent requests and had an average response rate of 244 ms.

#### Supported data types and public Xena Hubs

Today, cancer genomics research studies commonly collect data on somatic mutations, copy number, and gene expression, with other data types being relatively rare. However, as genomics technology advances, we expect these rarer data types to increase in frequency and new data types to be produced. With this in mind we designed Xena to be able to load any tabular or matrix formatted data, giving us exceptional flexibility in the types of data we can visualize, such as ATACseq peak signals (Supplemental Figure 1) and structural variant data (Supplemental Figure 10), a significant advantage over the UCSC Cancer Browser. Current supported data modalities include somatic and germline SNPs (Single Nucleotide Polymorphisms), INDELs, large structural variants, copy number variation, gene-, transcript-, exon-, protein-, miRNA-expression, DNA methylation, ATAC-seq peak signals, phenotypes, clinical data, and sample annotations.

UCSC Xena provides interactive online visualization of seminal cancer genomics datasets through multiple public Xena Hubs. We host over 1600 datasets from more than 50 cancer types, including the latest from TCGA, ICGC, TCGA Pan-Cancer Atlas (Hoadley 2018), and the GDC (Supplemental Table 1). Xena Hubs offer a significant and important performance advantage over these resources’ native APIs, especially when visualizing more than just a few samples. We use custom ETL (Extract-Transform-Load) processes to keep the Xena Hubs updated with the latest data from their respective sources (Supplemental Figure 1). We only download and process the derived datasets from each source, such as gene expression values, leaving the raw sequencing data at their respective locations. Xena complements each of these resources by providing powerful interactive visualizations for these data.

In addition to these well-known resources, we also host results from the UCSC Toil RNAseq recompute compendium, a uniformly re-aligned and re-called gene and transcript expression dataset for all TCGA, TARGET and GTEx samples (Vivian 2017). This dataset allows users to compare gene and transcript expression of TCGA ‘tumor’ samples to corresponding GTEx ‘normal’ samples. The UCSC Public hub hosts data curated from various publications.

### Conclusion

UCSC Xena complements existing tools including the cBioPortal, ICGC Portal, GDC Portal, IGV, and St. Jude Cloud (Ma 2018) in a number of ways. First, our focus is on providing researchers a lightweight, easy-to-install platform to visualize their own data as well as data from the public sphere. By visualizing data across multiple hubs simultaneously, Xena differentiates itself from other tools by enabling researchers to view their own data together with consortium data while still maintaining privacy. Further, Xena focuses on integrative visualization of multi-omics datasets across different genomic contexts, including genes, genomic elements, or any genomic region, for both coding and non-coding parts of the genome. Finally, Xena is built for performance. It can easily visualize of tens of thousands of samples in a few seconds and has been tested on single-cell data with up to a million cells. With single-cell technology, datasets will become orders of magnitude larger than traditional bulk tumor samples - Xena is well positioned to rise to this challenge.

While it is widely recognized that data sharing is key to advancing cancer research, how it is shared can impact the ease of data access. UCSC Xena is designed for cancer researchers both with and without computational expertise to easily share and access data. Users without a strong computational background can explore their own data by installing a Xena Hub on their personal computer using our installation and data upload wizards. Bioinformaticians can install a private or public Xena Hub on a server, in the cloud, or as part of an analysis pipeline, making generated data available in a user-friendly manner that requires little extra effort. Security for private hubs shared with a limited set of researchers is currently provided by protecting the computer itself such as using a firewall, in the future, we plan to develop hub-wide user authorization capability. This would be useful for collaborative projects who share private data with users across multiple institutions. It will also allow integration with existing federated authentication and authorization services. Data sharing has, and will continue to, advance cancer biology and Xena is part of the technological ecosystem that supports this.

UCSC Xena is a scalable solution to the rapidly expanding and decentralized cancer genomics data. Xena’s architecture, with its web-browser-based visualization and separate data hubs, allows new projects to easily add their data to the growing public compendium. We support many different data modalities, both now and in future, by maintaining flexible input formats. Xena excels at showing trends across cohorts of samples, cells, or cell lines. While we have focused on cancer genomics, the platform is general enough to host any functional genomics data. In this age of expanding data resources, Xena’s design supports the ongoing data sharing, integration, and visualization needs of the cancer research community.

## Acknowledgements

Research reported in this publication was supported by National Cancer Institute of the National Institutes of Health under award numbers 5U24CA180951-04 and 5U24CA210974-02. The content is solely the responsibility of the authors and does not necessarily represent the official views of the National Institutes of Health. This project has also been made possible in part by grant number 2018-182812 from the Chan Zuckerberg Initiative DAF, an advised fund of Silicon Valley Community Foundation. We would also like to thank AWS Cloud Credits for Research and Google Summer of Code.

