## supplementary materials for "The UCSC Xena platform for public and private cancer genomics data visualization and interpretation"

#### Supplemental Figure 1

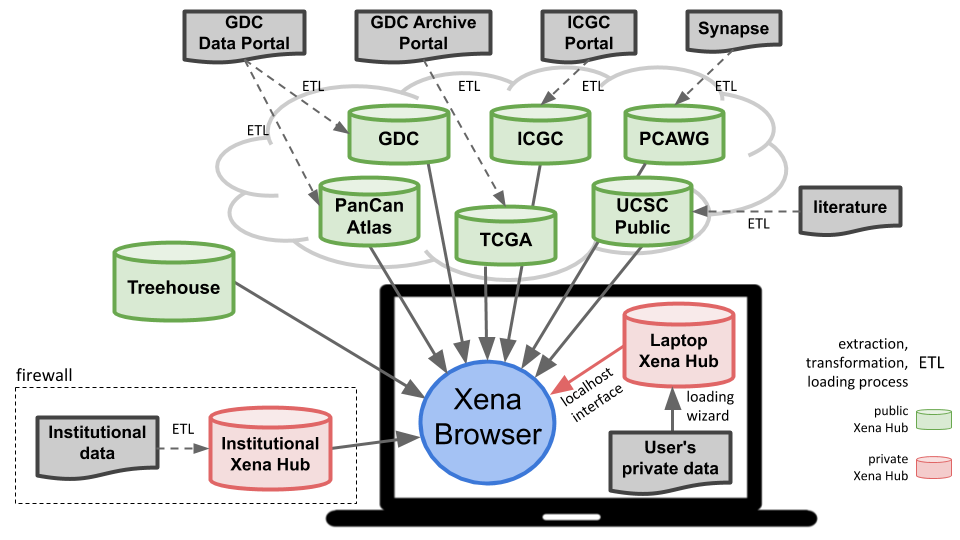

**Supplemental Figure 1.** Diagram of the UCSC Xena platform architecture. Multiple Xena Hubs (each shown as a database icon) are connected to the Xena Browser simultaneously. Data only flows from hub to browser. Public hubs are in green and private hubs in red. In this example, private data from an independent research collaboration (in red) is loaded into their own private Xena Hubs on their servers. Similarly, user's private data (in red) is loaded into a private Xena Hub on a researcher's computer. Data integration occurs within the Xena Browser on the user's computer. ETL (Extract-Transform-Load) processes bring data from outside sources into their respective public Xena Hubs. Xena Hubs offer a significant and important performance advantages over these resources’ native APIs, especially when visualizing many samples or cells. This design achieves data integration across both public and private resources while maintaining each hub’s data confidentiality.

#### Supplemental Figure 2

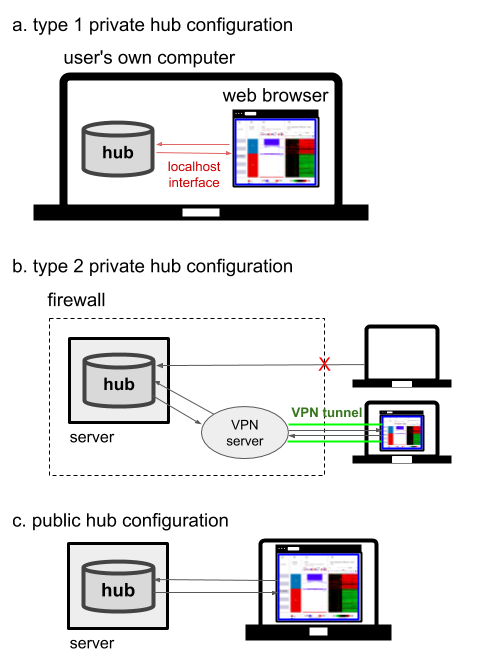

**Supplemental Figure 2.** Three types of Xena Hubs. **(a)** type 1 private hub configuration: hub is installed on a user’s own computer. It is configured to only respond to requests from the computer’s localhost network interface (i.e. http://127.0.0.1). This ensures that the hub only communicates with the computer on which the hub is installed. **(b)** type 2 private hub configuration: hub is installed on an outside computer. Hub is configured to respond to requests originating from external computers, however access to the computer is controlled via a firewall or similar technology. The hub uses the security provided by the computer to secure the data. Computers who request data without authorization will not have access. **(c)** public hub configuration: hub is installed on outside computer. Hub is configured to respond to requests originating from external computers and are not blocked by a firewall.

#### Supplemental Figure 3

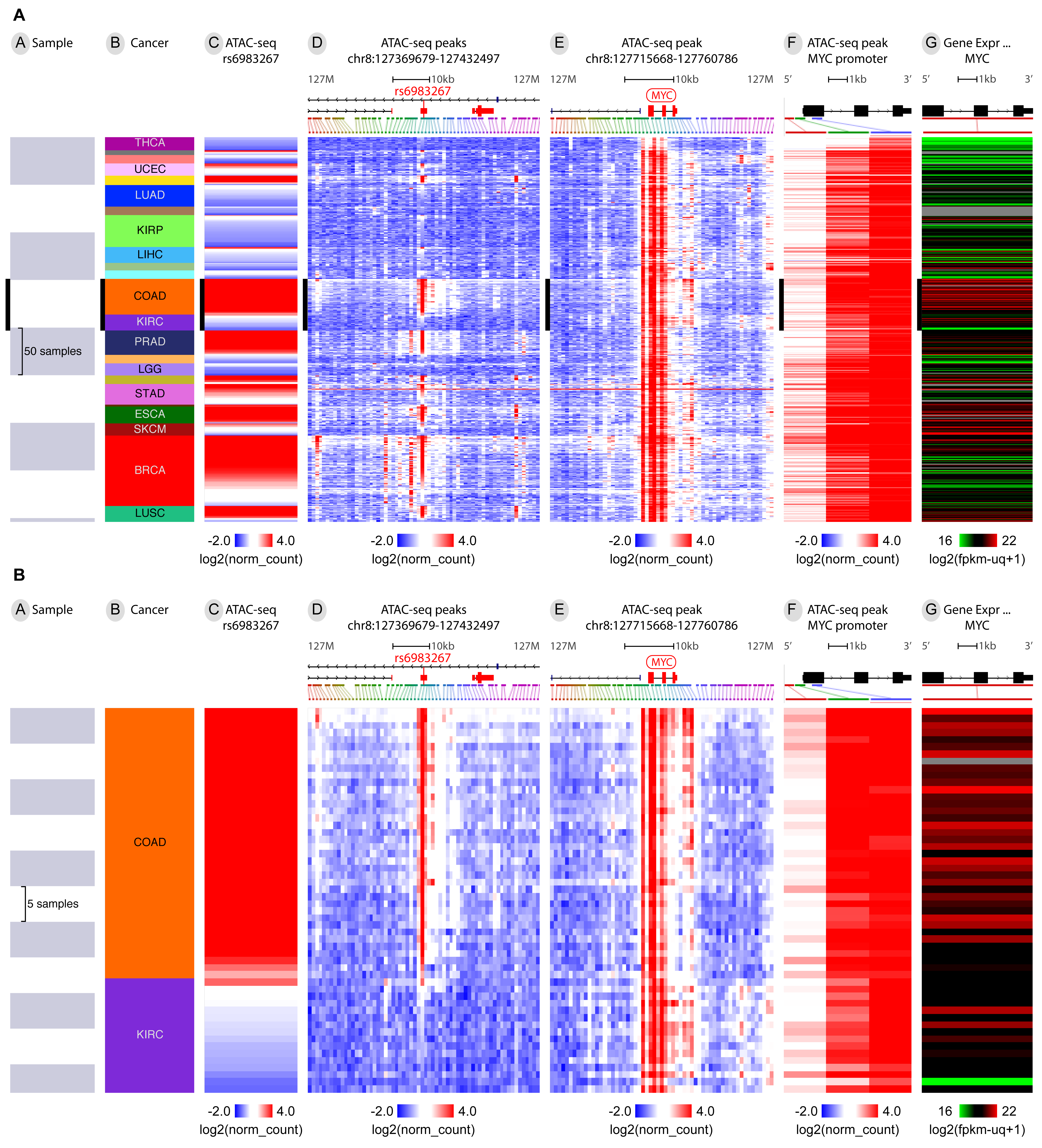

**Supplemental Figure 3. (a)** Xena Visual Spreadsheet examining the relationship between MYC gene expression (Column G), chromatin accessibility (ATAC-seq peaks in Column C through F, gene model at the top showing red to blue for increased to decreased chromatin accessibility), and cancer type (Column B) in TCGA. COAD (Colon adenocarcinoma) and KIRC (Kidney renal clear cell carcinoma) samples are highlighted with black bars across all columns using Xena’s dynamic find and highlight feature. Here we see that rs6983267, a functionally validated GWAS colon and prostate cancer susceptibility SNP, is more accessible in COAD, especially when compared to KIRC (Column C). Looking at the surrounding peaks (Column D) we can see that this pattern is localized to this SNP. Accessibility at this SNP is also associated with accessibility at MYC (Column E) and specifically the MYC promoter (Column F). Looking across all cancer types we see these COAD-specific findings may be applicable to other cancer types with extensive chromatin accessibility at 5’ and 3’ DNA elements, as suggested in Corces et al 2018. <https://xenabrowser.net/heatmap/?bookmark=032e821bfa82eb3c0f522877dbe3b252>

**(b)** Zooming in on COAD and KIRC we can more clearly see the contrast between these two cancer types. <https://xenabrowser.net/heatmap/?bookmark=471618c9813dabdee556bbaae43cf883>

#### Supplemental Figure 4

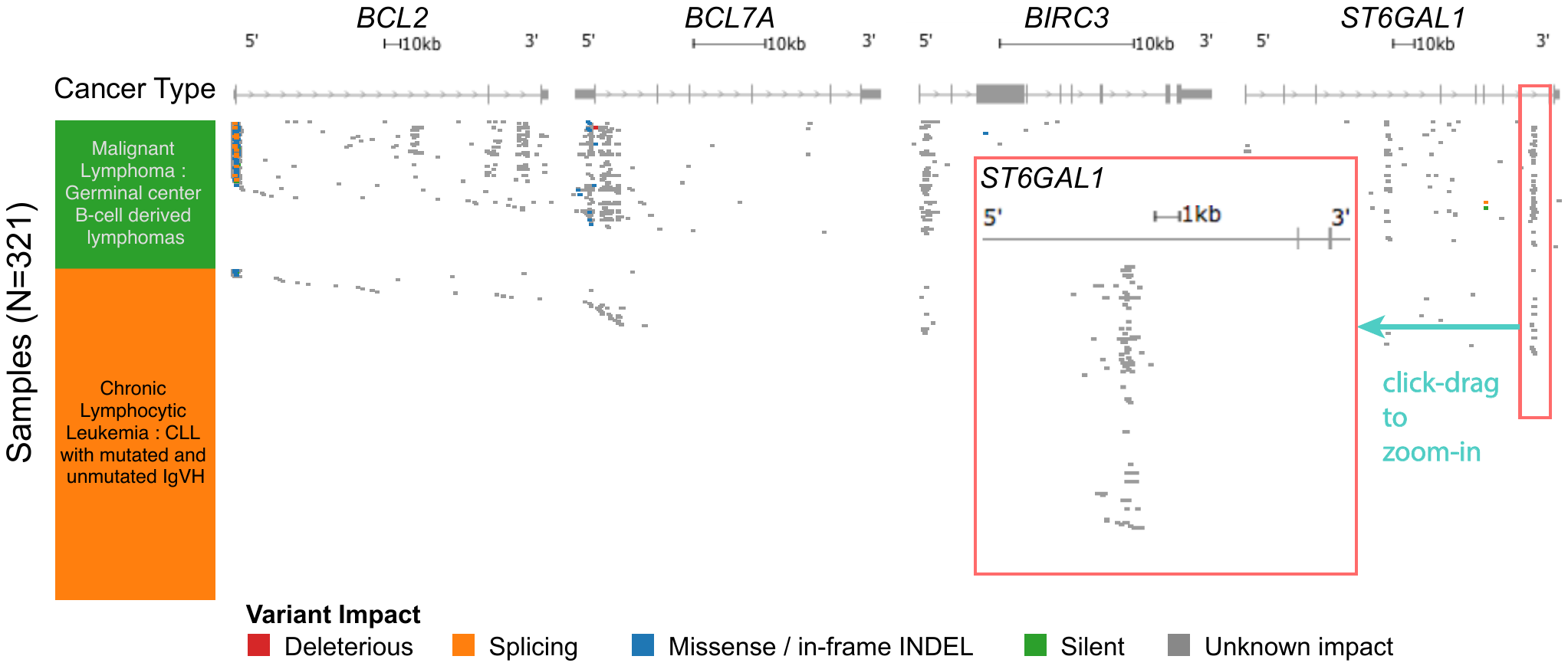

**Supplemental Figure 4.** Visualization of both coding and non-coding mutations from a gene-centric perspective in ICGC lymphomas. The columns left to right are cancer type, BCL2, BCL7A, BIRC3 and ST6GAL1 mutation status, respectively. Gene diagrams are shown at the top of each column, with exons as gray boxes, introns as lines. The position of each mutation is marked in relation to the gene diagram and colored by its functional impact. This figure shows the intronic mutations hotspots in these genes. These mutation 'pile-ups' would be not be visible if viewing exomes only. A dynamic toggle allows user to show or hide introns from the view. While the majority of the intronic mutations in this view have an unknown impact (shown in grey), they overlap with known enhancers regions (Mathelier 2015). Insert is a zoomed-in view of one of the hotspots in ST6GAL1. Users click-and-drag to zoom. <https://xenabrowser.net/heatmap/?bookmark=a11909d2c2c629ee999e1a9802fac7dd>

Supplemental Figure 5

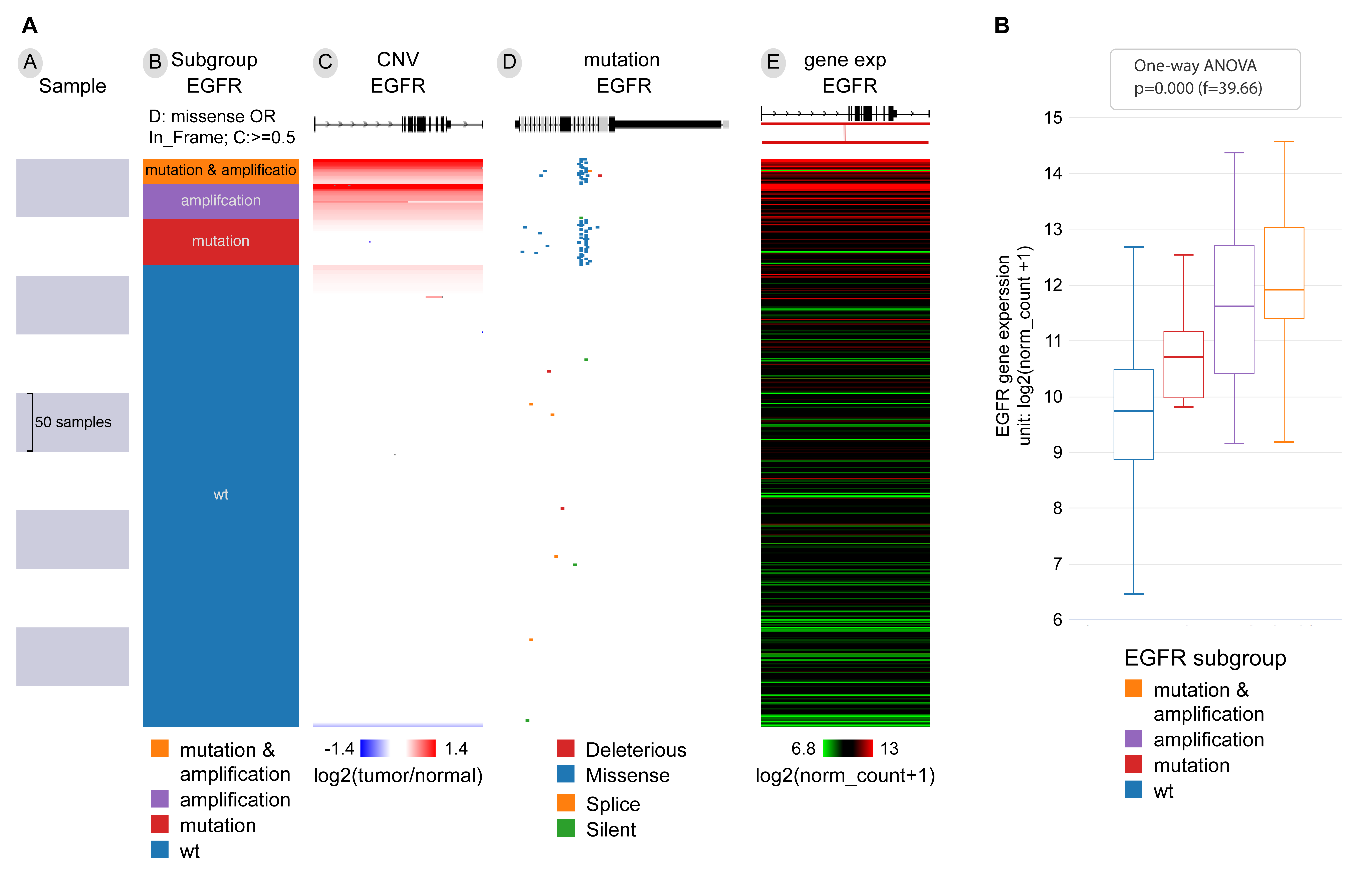

**Supplemental Figure 5. (A)** Xena Visual Spreadsheet showing EGFR status in TCGA Lung Adenocarcinoma. Column B is the EGFR subgroup where blue are EGFR wild-type samples, red are samples with an EGFR missense mutation or in frame deletion, purple are samples that have an amplification of EGFR (copy number value greater than 0.5), and orange are samples that have both an amplification and altered mutation status. Column C is EGFR copy number variation status, Column D is EGFR mutation status, and Column E is EGFR expression. This view shows that copy number amplification, mutation status, or both can drive high EGFR expression. <https://xenabrowser.net/?bookmark=5f3af5136b4ee90871da0b7fb81cb083>

(B) Here we can see the average EGFR expression for each of the 4 EGFR subgroups. Samples with both an amplification and altered mutation status have the highest gene expression. <https://xenabrowser.net/?bookmark=45b31e1e4fa268412abb7040c81057fb>

Supplemental Figure 6

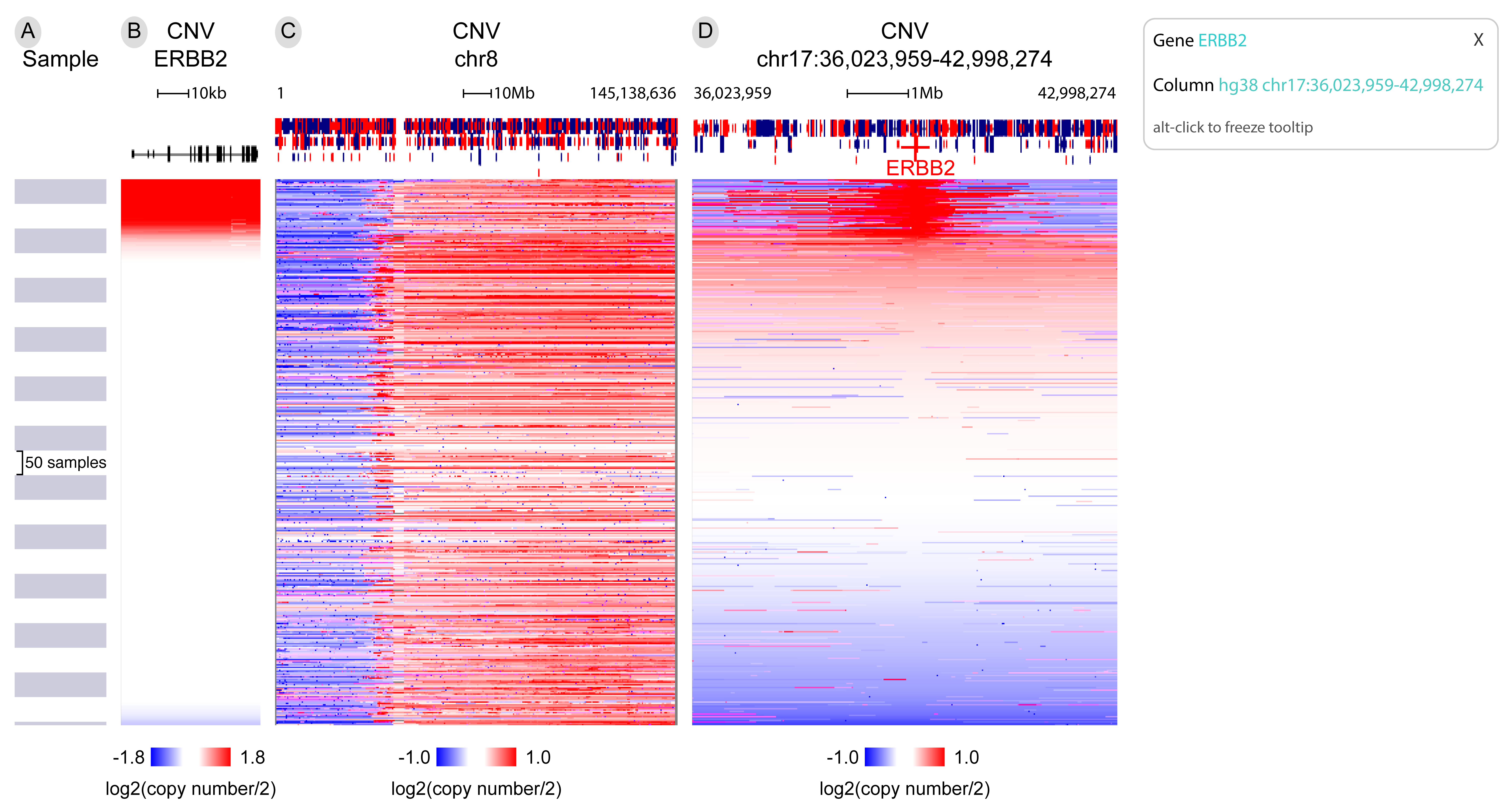

**Supplemental Figure 6.** A Xena Visual Spreadsheet contrasting gene-level and chromosome-arm-level copy number amplifications in TCGA Breast Cancer. Columns B through D show copy number status for just ERBB2, the entirety of chromosome 8, and ERBB2 and its flanking regions. We can see in column B that some samples have amplifications in ERBB2, but that this amplification is local to the ERBB2 gene (comparing Column B to Column D). In contrast we can see that many samples have an amplification of the entirety of the p-arm of chromosome 8, which is not localized to any one gene. In the upper right we can see an example of Xena’s tooltip, which shows more information about the data under the mouse cursor. <https://xenabrowser.net/?bookmark=9127f68a4b1547cad2bc493b97508416>

Supplemental Figure 7

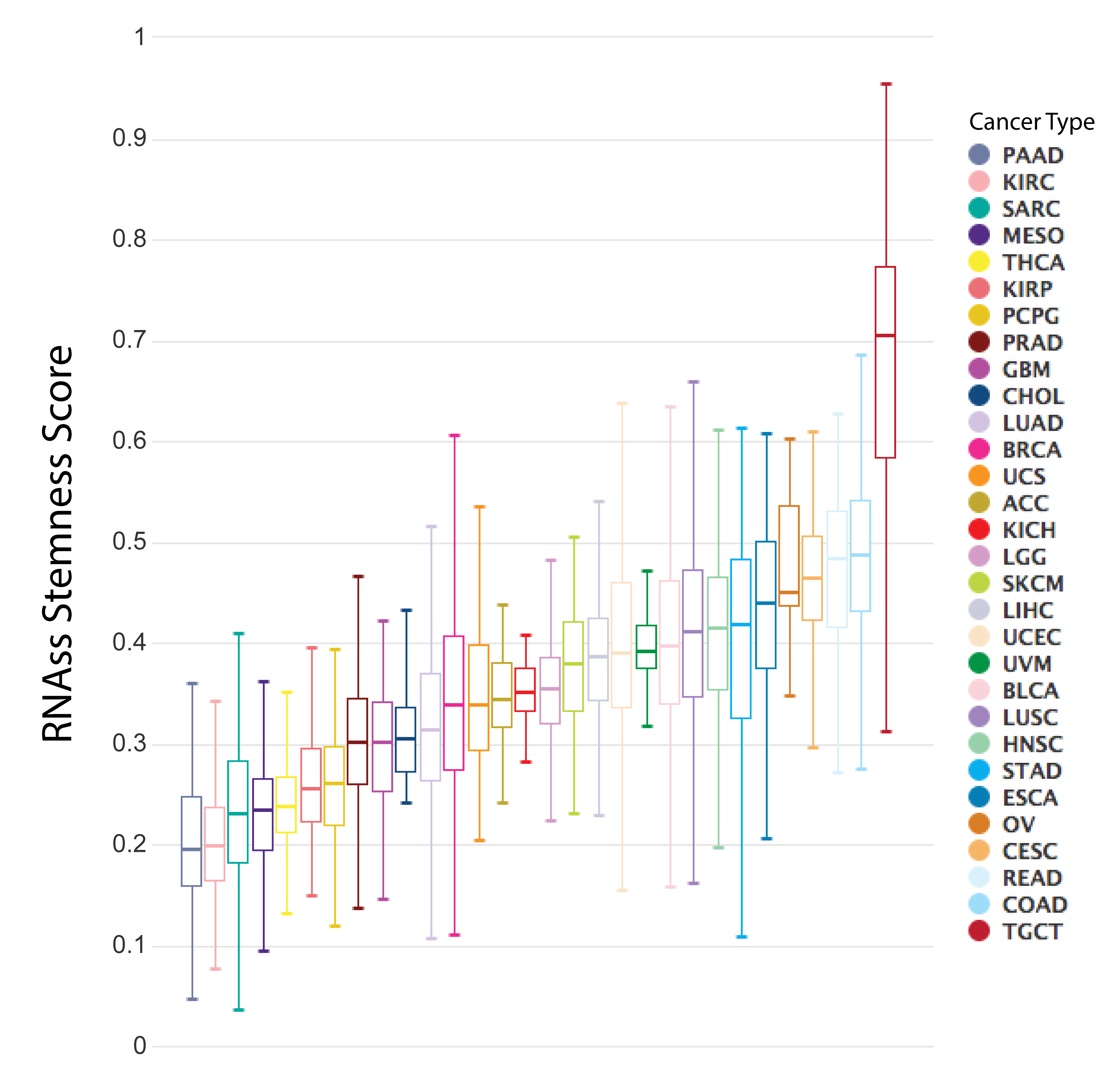

**Supplemental Figure 7.** The RNA-based stemness score as called by the PanCan Atlas Project across cancer type (Malta 2018). Cancers progress in part through the gradual gain of stem-cell-like features. Malta 2018 provides novel stemness indices for assessing this oncogenic de-differentiation. Here we can see that Testicular Germ Cell Tumors (TGCT) has a highest median stemness score compared to all other cancer types. <https://xenabrowser.net/?bookmark=7deeea053f221ba9fc506be855f353ac>

#### Supplemental Figure 8

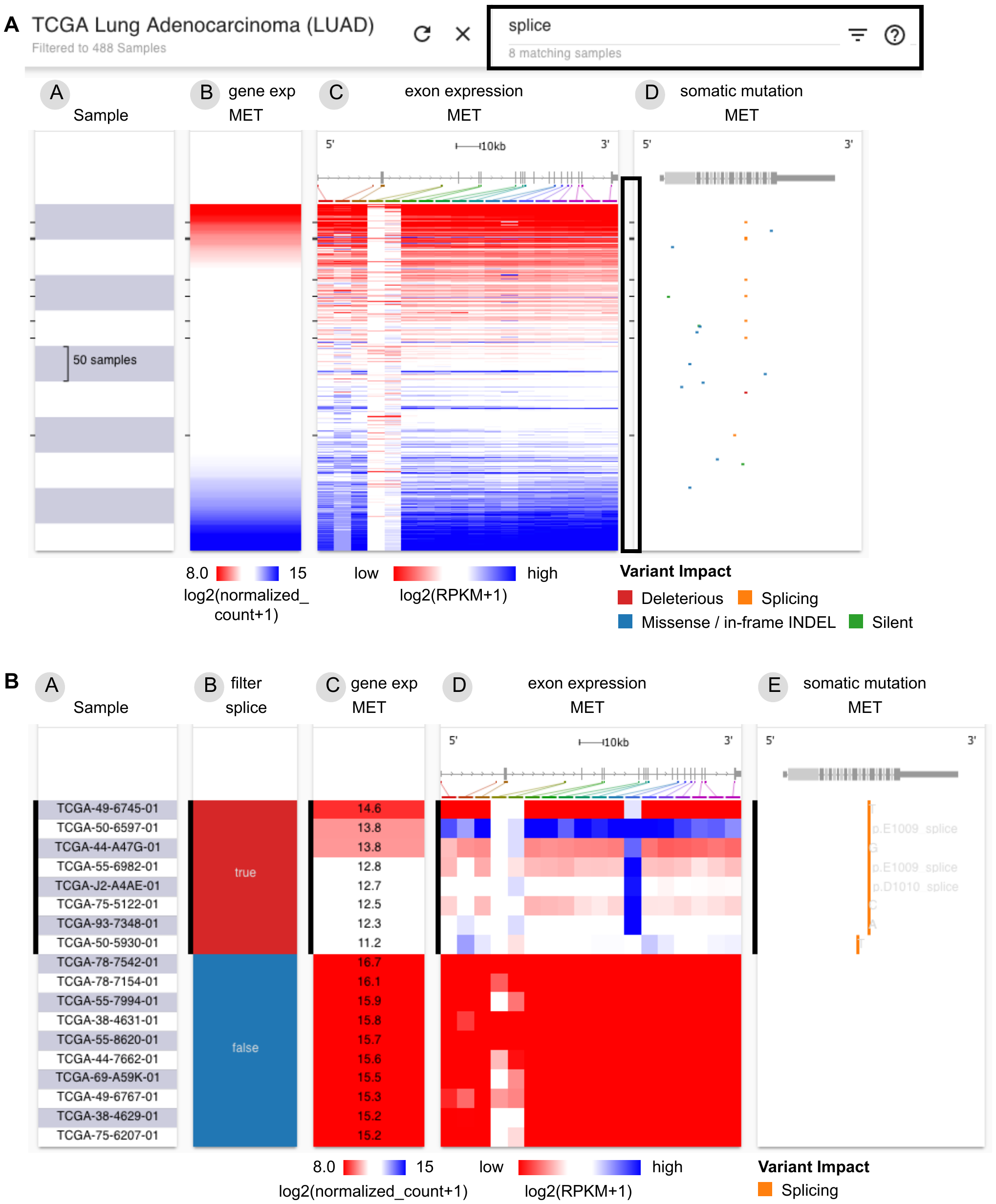

**Supplemental Figure 8.** Xena Browser text-based find, highlight, filter, and subgroup samples functionality. **(a)** Finding and highlighting samples in TCGA lung adenocarcinoma cohort that have a splice mutation in the gene *MET*. Similar to the ‘find in document’ feature in Microsoft Word, users can search all data on the screen. In this figure, the Xena Browser searched all columns for the user's search term 'splice' and highlighted samples with a 'splice' mutation with black tick marks (highlighted by the black box). More complex search terms can include 'AND', 'OR', '>', '<', and '='. Users can dynamically filter, zoom, and create subgroups based on the search results. Columns from left to right are MET gene expression, MET exon expression, and MET somatic mutation status. <https://xenabrowser.net/heatmap/?bookmark=c5873fb094ef714e44e65df217e93071>

**(b)** After creating a new column with two subgroups. Columns are same as (a) with the user-generated column inserted on the left. Samples that matched the query of 'splice' were assigned a value of "true" and those that do not "false". The researcher has zoomed to a few samples at the top for a more detailed view. The figure shows that samples that have the splice site mutation (orange tick marks, column E) have lower expression of MET’s exon (column D). The splice mutation causes exon 14 skipping and results in the activation of MET (Kong-Beltran 2006, The Cancer Genome Atlas Research Network 2014). <https://xenabrowser.net/heatmap/?bookmark=748c42b0b49552004da53873950aad62>

#### Supplemental Figure 9

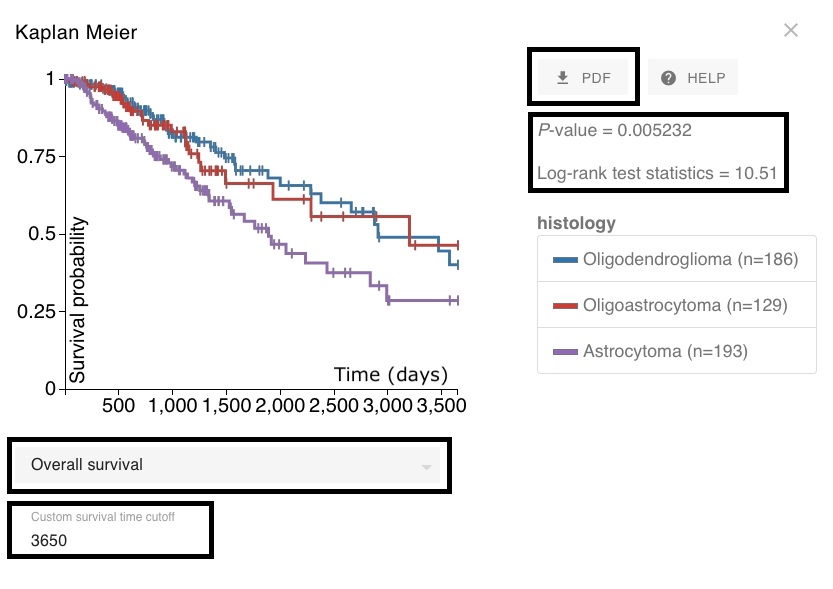

**Supplemental Figure 9.** Kaplan-Meier analysis of overall survival for TCGA lower grade glioma histological subtypes. Black boxes in the figure highlight, top to bottom, a button to generate a PDF, the statistical analysis results, a dropdown menu to select different survival endpoints such as overall or recurrence-free survival, and a textbox to enter a custom survival time cutoff (currently set to 3,650 days, or 10 years). This figure shows that patients characterized as having the astrocytoma histological subtype have significantly worse 10-year overall survival compared to the oligodendroglioma and oligoastrocytoma subtypes (p < 0.05). <https://xenabrowser.net/heatmap/?bookmark=2f9d783982879594dd0f52564058372d>

#### Supplemental Figure 10

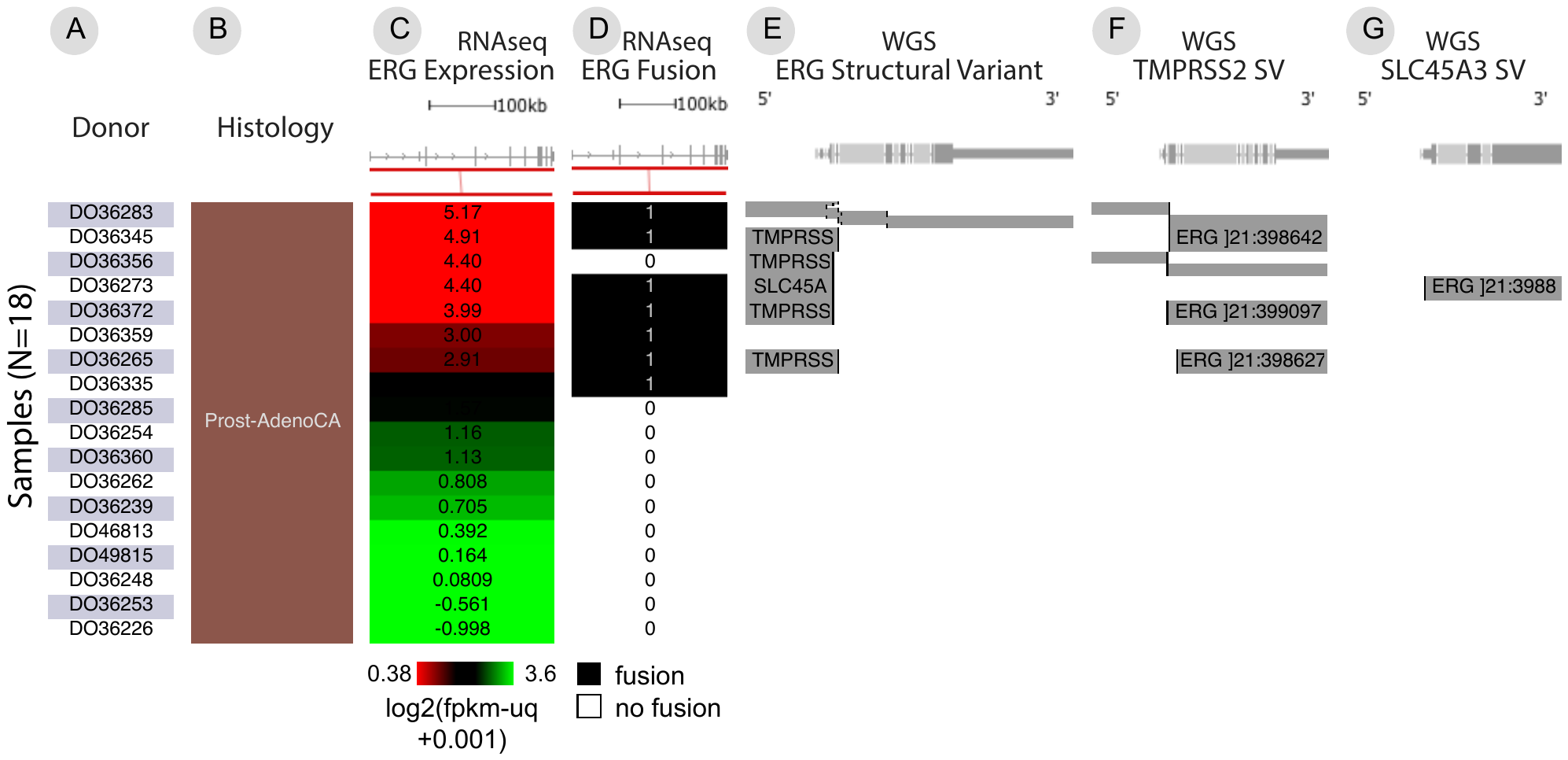

**Supplemental Figure 10.** Visualization of large structural variants. This figure shows frequent ERG fusion in PCAWG prostate cancer detected by both RNA-seq and DNA-seq analysis. The left three data columns (B, C, D) are histology, ERG gene expression, and ERG fusion detected using RNA-seq data. In the ERG fusion column (D), samples that have a fusion are marked with 1 and those that do not are marked with 0. The next three columns (E, F, G) show structural variant calls made using whole-genome DNA-seq data for ERG, TMPRSS2, and SLC45A3. Precise breakpoints are mapped to gene diagrams. A grey bar indicates an external piece of DNA that is fused at the breakpoint. Gene names on the grey bars show the origin of the external DNA that is joined. This figure shows that TMPRSS2 and SLC45A3 are fusion partners for ERG, and that these fusions correlate with over-expression of ERG. Fusions detected by RNA-seq and whole-genome sequencing are not always consistent. Here, even using a consensus of DNA-based detection methods, one fusion detected by a consensus of RNA-based detectors is missed, and the converse is also seen. This example highlights that an integrated visualization across multiple data types and algorithms provides a more accurate model of a genomic event. <https://xenabrowser.net/heatmap/?bookmark=92db485580786d1ef14c6c06b680201b>

#### Supplemental Table 1

| **Public Xena Hub Name and URL address** | **Samples** | **Cohorts** | **Data Content** |
| --- | --- | --- | --- |
| TCGA  https://tcga.xenahubs.net | 12,839 | 38 | TCGA copy number, gene-, exon-, miRNA-, and protein-expression, somatic mutation, DNA methylation, survival, and clinical data |
| Pan-Cancer Atlas  https://pancanatlas.xenahubs.net | 12,839 | 1 | TCGA copy number, gene-, miRNA-, and protein-expression, somatic mutation, DNA methylation, molecular subtypes, curated survival data |
| GDC  https://gdc.xenahubs.net | 19,189 | 41 | TCGA and TARGET copy number, somatic mutations, gene- and miRNA-expression, DNA methylation, overall survival, and clinical data |
| ICGC  https://icgc.xenahubs.net | 17,697 | 4 | ICGC copy number, gene expression, somatic mutation |
| PCAWG  https://pcawg.xenahubs.net | 2,834 | 2 | whole-genome copy number, somatic mutations, large structural variants, gene- and miRNA-expression, purity, ploidy, mutational signatures, survival, and curated histology |
| UCSC Toil RNAseq  https://toil.xenahubs.net | 19,131 | 5 | TCGA, TARGET, and GTEx gene- and transcript-expression |
| ATAC-seq  https://atacseq.xenahubs.net | 404 | 2 | TCGA ATAC-seq peak signal |
| UCSC Public  https://ucscpublic.xenahubs.net | 21,486 | 36 | copy number, gene expression, mutation, survival, drug response, and clinical data from various publications |

**Supplemental Table 1.** Summary of data hosted on public Xena Hubs as of September 6, 2019.

### Supplemental NoteMethods

#### Xena’s privacy aware architecture

Xena does not utilize a central rendering service or require hubs to be publicly accessible on the internet like, for example, the UCSC Genome Browser does. Data flows in one direction, from the Xena Hubs to the Xena Browser. If the user installs a Xena Hub on their laptop, the hub is as secure as the laptop. Xena Hubs are by default configured to only accept data requests coming from the localhost network interface (i.e. loopback device). If the user installs a Xena Hub on a local network behind a firewall, the hub is as secure as the local network. These Xena Hubs can be configured to bind an external port, making the data accessible to the local network. Users can do this by binding an external interface by using the IP associated with that interface or using 0.0.0.0 to bind all interfaces.

The Xena Browser accesses data from a local Xena Hub on the same computer by requesting data from <http://127.0.0.1>. The local Xena Hub will make the data within it available at this address. The local Xena Hub will only answer requests made form the user's own computer. At the time of writing, this functionality is only supported by some web browsers. This includes Chrome, and Firefox, but not Safari.

Note that a very limited set of metadata is considered to be not secure in the Xena model. This includes cohort IDs and samples IDs. This metadata is visible to other hubs in the following scenarios. When the user selects a cohort, all hubs are queried for samples on that cohort. When the user selects a data field, the hub holding that field is queried with all the sample IDs in the cohort. This means, for example, that two hubs holding data on the same cohort will see the union of sample IDs from that cohort. While data queries are not made available publicly, the user who set up the hub can comb through logs for these queries. For these reasons, these metadata fields should not contain private information.

#### Xena Hub

The Xena Hub is a JVM-based application, written in Clojure, that serves functional genomic data over HTTP. It exposes a relational query API for data slicing and metadata. We decided to use a query language instead of REST for our APIs because it allowed us to decouple the client and server. To support interactive visualization, REST APIs would have to be denormalized for performance (e.g. by joining related objects, and projecting the result). This would require a tight coupling between the REST endpoints and particular views, and therefore frequent server redeployment to match browser client updates. Using REST becomes impractical because all the hubs in the Xena System will need to be updated including the ones people installed on their laptops or servers. In contrast to REST APIs, a query language allows us to fetch exactly the data we need, and only the data we need, for quickly evolving visualizations and data shapes, without redeployment of the hubs. This is similar in motivation to Facebook's GraphQL, and Netflix's Falcor, but was developed before these libraries were available.

Internally, Xena Hubs use the H2 database for storage. Data is stored in opaque blocks in a column orientation, which allows fast retrieval of a field for all samples of a dataset, or a subset of samples. A hub can be installed either via the command line or via the point-and-click install4j graphical user interface.

#### Xena Browser

The Xena Browser is a javascript application to visualize and analyze functional genomics data stored in one or more Xena Hubs (Langmead 2018, Cieślik 2018). We support modern web browsers including Chrome, Firefox and Safari. The primary technologies are React, the 2D canvas API, and RxJS. Babel is used for es6 support, and webpack for the build. The application architecture is an asynchronous model similar to redux-observable (<https://redux-observable.js.org/>), with semantic actions that update application state, and action side-effects creating Rx streams that will dispatch later actions. The redux (<https://redux.js.org/>), or Om (<https://github.com/omcljs/om>), pattern of immutable, single-atom state makes it simple to keep multiple views in sync, and provides “time travel” debugging during development.

We prefer the canvas API to SVG libraries such as D3, because it performs better at large data scales. With the advances in javascript JIT compilers, we find that optimized loops over canvas pixel buffers out-perform geometric drawing primitives, such as *rect()*, and *stroke()*, when rendering dense views of large data.

The jsverify property-based testing library (<http://jsverify.github.io/>) is used for unit and integration testing. Property-based, or "generative" testing is similar to fuzzing -- generating random test cases, and asserting invariants over the results -- but on failure, attempts to find a minimal failing test case. This usually results in more tractable failure cases. Property-based testing allows us to test a much larger portion of the input space than conventional "known-answer" unit tests, and frequently identifies failure cases that we would never think to test.

All of our code is open source and available for reuse under and Apache 2.0 license (Xena Hub: <https://github.com/ucscXena/ucsc-xena-server>; Xena Browser: <https://github.com/ucscXena/ucsc-xena-client>; all other code: <https://github.com/ucscXena>). We also have contributed two javascript modules to BioJS (Gómez 2013), including a Kaplan-Meier module (<https://github.com/ucscXena/kaplan-meier>) to compute Kaplan-Meier statistics, and a static-interval-tree library (<https://github.com/ucscXena/static-interval-tree>) to effectively find overlapping intervals in one dimension.

#### Public Xena Hubs

We download functional genomics data from each respective source: GDC data portal (<https://portal.gdc.cancer.gov/repository>) for the GDC Hub, GDC legacy archive (<https://portal.gdc.cancer.gov/legacy-archive>) for the TCGA Hub, ICGC data portal (<https://dcc.icgc.org/>) for the ICGC Hub, Synapse TCGA Pancan Atlas Data project (<https://www.synapse.org/#!Synapse:syn3241074/wiki/194741>) for the Pan-Cancer Atlas Hub, Synapse ICGC-TCGA Whole Genome Pan-Cancer Analysis project (<https://www.synapse.org/#!Synapse:syn2351328/wiki/62351>) for the PCAWG hub, Vivian et. al. 2007 for the UCSC RNAseq Recompute Toil Hub, and for the GDC’s Publications site (<https://gdc.cancer.gov/about-data/publications/ATACseq-AWG)> for ATACseq hub. We also host data curated from various publications such as CCLE (Cancer Cell Line Encyclopedia, Barretina 2012) and MET500 (Robinson 2017) in the UCSC Public Hub. The GDC, ICGC and UCSC Public hubs are updated periodically.

Using custom ETL processes, the downloaded data were wrangled into a generic tabular or matrix format and loaded into the corresponding Xena Hubs. Specific wrangling steps, including any normalization, is listed for each dataset in the Xena Browser dataset pages (<https://xenabrowser.net/datapages/>). The wrangled data is available for bulk download from the dataset pages. Approximately 1TB of data are downloaded each month from our hubs. For example, 1.5 TB of compressed data were downloaded in the month of July 2019. We also offer programmatic access to slices of data through the Xena python package (<https://github.com/ucscXena/xenaPython>), which can be used independently or in a Jupyter Notebook. The independently developed UCSCXenaTools R package (Wang 2019) offers additional programmatic access in R.

We deploy all public-facing Xena Hubs in the Amazon Web Services (AWS) cloud-computing environment. Each hub is built using an AWS elastic load balancer connected to two EC2 r5.4xlarge servers. This architecture ensures fast performance even when user queries are highly concurrent. Response rates in this environment are, on average, 244 ms with 50 users making concurrent requests (tested using the TCGA Breast Cancer cohort data).

Treehouse Hub

An example of a public hub hosted by an institution is the Treehouse Hub ([https://xena.treehouse.gi.ucsc.edu](https://xena.treehouse.gi.ucsc.edu/)) set up by the set up by the Childhood Cancer Initiative (<https://treehousegenomics.soe.ucsc.edu)>. It hosts RNAseq gene expression data of pediatric cancer samples provided by Treehouse's clinical partners and repositories, harmonized with the UCSC RNAseq recompute compendium dataset of TCGA, TARGET (Therapeutically Applicable Research To Generate Effective Treatments) and GTEx (Genotype-Tissue Expression, GTEx Consortium 2017) samples (Morozova 2017, Vivian 2017). This data is used to facilitate interpretation of a pediatric sample in the context of a large pan-cancer cohort, as the same bioinformatics pipeline processed all samples. Since the data hub is set up and hosted separately, The Treehouse project has complete control over the data and data access since they set up and host the Xena Hub themselves. The UCSC Xena platform allows them to host and update their public data, allowing it to be downloaded and visualized.

#### User-centered design principles

The Xena System was developed using User Experience Design methodologies. User-Centered Design is design based upon an explicit understanding of users, tasks, and environments, and is driven and refined by iterative user-centered evaluation. We use need-finding interviews, prototypes, wireframes, and user acceptance testing to help ensure that Xena meets user needs.

Data availability statement

No datasets or data were analyzed during this study
